# Zebrafish sp7 mutants show tooth cycling independent of attachment, eruption and poor differentiation of teeth

**DOI:** 10.1101/259085

**Authors:** E Kague, P.E Witten, M Soenens, CL Campos, T Lubiana, S Fisher, C Hammond, Brown K Robson, MR Passos-Bueno, A Huysseune

## Abstract

The capacity to fully replace teeth continuously makes zebrafish an attractive model to explore regeneration and tooth development. The requirement of attachment bone for the appearance of replacement teeth has been hypothesized but not yet investigated. The transcription factor *sp7* (*osterix*) is known in mammals to play an important role during odontoblast differentiation and root formation. Here we study tooth replacement in the absence of attachment bone using *sp7* zebrafish mutants. We analysed the pattern of tooth replacement at different stages of development and demonstrated that in zebrafish lacking *sp7*, attachment bone is never present, independent of the stage of tooth development or fish age, yet replacement is not interrupted. Without bone of attachment we observed abnormal orientation of teeth, and abnormal connection of pulp cavities of predecessor and replacement teeth. Mutants lacking *sp7* show arrested dentinogenesis, with non-polarization of odontoblasts and only a thin layer of dentin deposited. Osteoclast activity was observed in *sp7* mutants; due to the lack of bone of attachment, remodelling was diminished but nevertheless present along the pharyngeal bone. We conclude that tooth replacement is ongoing in the *sp7* mutant despite poor differentiation and defective attachment. Without bone of attachment tooth orientation and pulp organization are compromised.

## Introduction

Considerable progress in understanding tooth development and the molecular machinery that controls it has been achieved over the past decades (Balic and Thesleff, 2015). However, key aspects such as the capacity of full tooth regeneration in some vertebrates require better understanding to be able to translate such knowledge to the dental clinic. Like in other non-mammalian vertebrates, the dentition in zebrafish *(Danio rerio)* is replaced continuously throughout life (polyphyodont), with every tooth being replaced every 8–12 days (in juveniles). This makes zebrafish an attractive model to study odontogenesis, tooth replacement and regeneration (Huysseune, 2005; Stock, 2007; Witten et al., 2017).

Briefly, the zebrafish dentition is fully established around 26 days and consists of eleven teeth positioned on each fifth ceratobranchial (constituting the pharyngeal jaws). The teeth are arranged in three rows: ventral, mediodorsal and dorsal (Borday-Birraux et al., 2006; Van der heyden and Huysseune, 2000). The first tooth starts to develop at 48 hours post-fertilization (hpf) and at around 80 hpf attaches to the ceratobranchial, thereby becoming functional. Concomitantly, the first replacement tooth forms at the same position from the successional lamina, attached to the epithelial crypt of the predecessor tooth. The successional lamina thickens in the early morphogenesis stage, re-invaginates, and forms a bi-layered bell-shaped structure, consisting of outer and inner epithelial layer, during late morphogenesis stage (Borday-Birraux et al., 2006). Differentiation of pre-ameloblasts and pre-odontoblasts with deposition of the enameloid (a mixed epithelial-mesenchymal product) and dentin matrix at the tooth tip marks the cytodifferentiation stage. Beyond the tooth base, easily identified by the position of the epithelial cervical loop, attachment bone is deposited. The attachment bone eventually ankyloses the dentin base to the underlying ceratobranchial bone (Huysseune et al., 1998). Unlike the situation in mammals, there is no distinction between crown and root dentin, nor is there cementum. Attachment coincides with eruption (Huysseune and Sire, 2004) and marks the start of development of the replacement tooth. While the predecessor functional tooth is attached to the pharyngeal arch, the replacement tooth does not have any skeletal support. The role, if any, of tooth attachment to the underlying bone for initiation of the replacement tooth formation remains unclear. It has been suggested that attachment and eruption may trigger the formation of the replacement tooth. This hypothesis is based on the observed coincidence of eruption of the predecessor and formation of its successor (Borday-Birraux et al., 2006). It has also been suggested that eruption may lead to displacement and activation of a putative stem cell niche (Huysseune and Thesleff, 2004). However, to date this hypothesis has not been fully tested.

In mammals, root dentinogenesis and cementogenesis, i.e. the deposition of the root dentin and cementum matrix, respectively, rely on a cascade of signalling molecules that act sequentially, among them the transcription factor *Sp7/Osterix. Sp7* is required both for osteogenesis and odontogenesis (He et al., 2016; Sinha and Zhou, 2013). The gene has been associated with low bone mineral density from human genome wide association studies (GWAS) (Rivadeneira et al., 2009). Recently, a mutation was identified in a human patient with recessive osteogenesis imperfecta, in which along with poorly mineralized bones, tooth eruption was delayed (Lapunzina et al., 2010). *Sp7* knockout mice die prematurely after birth and lack most of the skeleton. Yet, despite arrest in odontogenesis and defective alveolar bone formation, normal tooth number and morphogenesis were observed, demonstrating the independence of (at least early stages of) tooth formation from tooth attachment (Nakashima et al., 2002). Using conditional knockout mice where *sp7* is deleted under the control of *col1a1* or *osteocalcin* promoters, hypoplastic dentin and root abnormalities were observed (Kim et al., 2015).

The importance of *sp7* in osteogenesis and dentinogenesis extends to other vertebrates besides mammals and the pattern appears to be conserved. In teleost fish, the role of *sp7* in bone and teeth has been recently reported (Kague et al., 2016; Niu et al;, 2017; Yu et al., 2017; Zhong et al., 2016). We have described previously that zebrafish null for *sp7* show severe defects in bone growth and mineralization, consistent with the roles ascribed to the gene through GWAS in humans, mouse and medaka fish. Moreover, mineralization of teeth was limited to their tips (Kague et al., 2016). In medaka, similar observations were made; the number and patterning of teeth appeared normal in mutants, although tooth sizes were generally smaller (Yu et al., 2017).

Here we describe lack of attachment bone in zebrafish lacking *sp7* and its role in tooth development and cycling. We show that *sp7* is expressed in differentiated odontoblasts of young, but not mature teeth. We show that *sp7-/-* mutants can achieve normal tooth patterning, yet teeth display abnormal orientations and dentinogenesis arrested at an early stage, with only a thin layer of dentin deposited. Tooth attachment does not occur, irrespective of developmental stage or age. Importantly, tooth replacement is not affected, showing that formation of the replacement tooth is autonomous from attachment, eruption or proper orientation of the predecessor tooth. Finally, our findings suggest a role of *sp7* in pulp mesenchyme organization.

## Material and Methods

### Zebrafish husbandry and lines

Zebrafish were raised and maintained under standard conditions (Westerfield, 2000). Zebrafish sp7^hu2790^ mutants were generated via target–selected ENU mutagenesis at the Hubrecht Institute (Wienholds et al., 2003), and maintained as described (Kague et al., 2016).

### Alizarin Red S Staining

Alizarin Red S staining was performed in fixed fish to label calcified tissues and carried out using standard protocols (Walker and Kimmel, 2007). Pharyngeal bone was cautiously dissected for whole mount imaging. Images were acquired on an Olympus SZX16 microscope with a DP72 camera. Live Alizarin Red S staining was carried out as previously described (Bensimon-Brito et al., 2016).

### Micro-Computed Tomography (µCT)

Adult fish, 1.5 year old, were fixed in 4% PFA followed by ethanol dehydration. Fish were scanned using SkyScan 1272 high resolution micro-CT scanner (Bruker) under voxel size of 5µm, x-ray source of 50kV and 200A and 0.5mm aluminium filter. Images were reconstructed using SkyScan CTAn software. Amira 6.0 was used for 3D volume rendering and image acquisition.

### Confocal microscopy

Transgenic fish *Tg(RUNX2:egfp)* (Kague et al., 2012) and *Tg(sp7:mcherry)* (Kague et al., 2016) were used to visualize osteoblasts/odontoblasts and *sp7* expression, respectively. Transgenic fish were live stained with Alizarin Red S overnight followed by dissection of the pharyngeal bone region. Samples were mounted in 1% LMP agarose (Invitrogen #16520050) followed by imaging under stereomicroscope (Leica msv269 and ddc700t camera) and Leica SP5II confocal microscopy (Leica LAS software) using 10x PL APO CS lens. Amira 6.0 was used for 3D rendering, segmentation and picture acquisition. For *sp7* expression *3D* volume renders were visualized using the Physics color for *sp7* and white for bone.

### Histology

*sp7* mutant fish of 6 weeks, 6 months and 1 year, along with wt siblings, were fixed in 4% PFA, decalcified in 50mM EDTA and dehydrated prior to embedding in Technovit. Serial sections (2-4 µm) were prepared using a Prosan HM360 microtome. A total of five mutant fish was serially sectioned, with ages ranging from 6 weeks to 6 months (standard length, S.L., between 9 and 18 mm). For comparison, we used serial sections of wt size-matched fish (used in (Huysseune, 2006); additionally, three adult wt siblings (S.L. 24 to 27 mm) were serially sectioned. Sections were stained with toluidine blue, dried and mounted using Depex. Observations were performed using a Zeiss Axio Imager Z1 microscope with a 5 Megapixel CCD camera.

### Tartrate-resistant acid phosphatase (TRAP) staining

TRAP staining to detect osteoclast activity was carried out on 5µm GMA (glycol methacrylate) sections following the protocol as previously described (Witten et al., 2001). Sections were incubated with naphthol AS-TR phosphate as enzyme substrate and hexazotized pararosaniline as a color component in acetate buffer (pH 5.5) containing 50 mM L(+) di-sodium tartrate dihydrate. Sections were counterstained with Mayers acidic hematoxylin.

### Immunohistochemistry

Zebrafish carrying the transgene *Tg(sp7:mcherry)* were used to evaluate expression of *sp7* (Kague et al., 2016). Pharyngeal jaws of adult fish were dissected, followed by imaging using fluorescent microscopy (Olympus SZX16 microscopy with DP72 camera). Immunohistochemistry was performed on cryosections. A primary DS-red rabbit polyclonal antibody (Clontech #632496) was used at 1:200 dilution and secondary goat anti-rabbit Dylight-488 antibody (Abcam AB98507) at 1:300 dilution, following a previously described protocol (Knopf et al., 2011).

## Results

### sp7 mutants lack tooth attachment and display severe mineralization defects

We have previously described overall delayed mineralization in zebrafish homozygous *sp7* mutants (sp7^hu2790-/-^). These mutants express a truncated sp7 protein lacking the three zinc finger domains required for DNA binding and are predicted to be null mutants (Kague et al., 2016). We have shown tooth tips with limited mineralization, hence failing to form a mineralized attachment to the pharyngeal bone. To investigate whether the attachment is defective throughout development, we carried out a morphological analysis of tooth mineralization at different developmental stages, from 3 weeks to 1.5 years, using Alizarin Red S staining.

Wild type fish (wt) at 3 weeks post-fertilization (wpf) showed clear and even Alizarin Red S staining in the pharyngeal bone and teeth (Fig. 1A, A’). In *sp7* mutants, the pharyngeal bone, although mineralized, showed unstained regions, reflecting delayed and/or poor mineralization. In mutants teeth possessed mineralized tips only; mineralization was absent along the tooth shaft, and no attachment was observed (Figure 1B, B’). The loss of Alizarin Red S staining after hypermineralization of the enameloid cap, normally seen in wt teeth, was not observed in the mutant dentition at any of the stages examined (Fig.1. C-F’). While thickening of the bones was observed during the ongoing development and growth of the zebrafish, attachment of the teeth to the pharyngeal bone in juvenile or adult mutant fish was not observed (Fig. 1D, D’, F, F’). In mutants, the shape of the pharyngeal bone differed from that seen in wt fish, showing a thinner structure with uneven staining (Fig.1 E, F, F’). To check if attachment would be observed in adult fish, we carried out micro-computed tomography (µCT) in 1.5 year old fish (Fig. 1G-H). Despite increased thickness of the pharyngeal bone, its shape remained as observed in young adults, and attachment of teeth was absent. The partial mineralization of the teeth gave an appearance of floating dots close to the pharyngeal bone. The phenotype characterized by the lack of attachment bone is fully penetrant, however phenotypic variability was observed among the mutants. Pharyngeal bone mineralization was completely lacking in severe cases while other specimens showed only mild mineralization defects. This variability was also observed in other parts of the skeleton, such as skull bones, as previously described (Kague et al., 2016). Thus, our data revealed that the *sp7* mutant has teeth with mineralized tips only and does not show tooth attachment to the bone in any stage of development or age of the fish.

**Fig 1.**
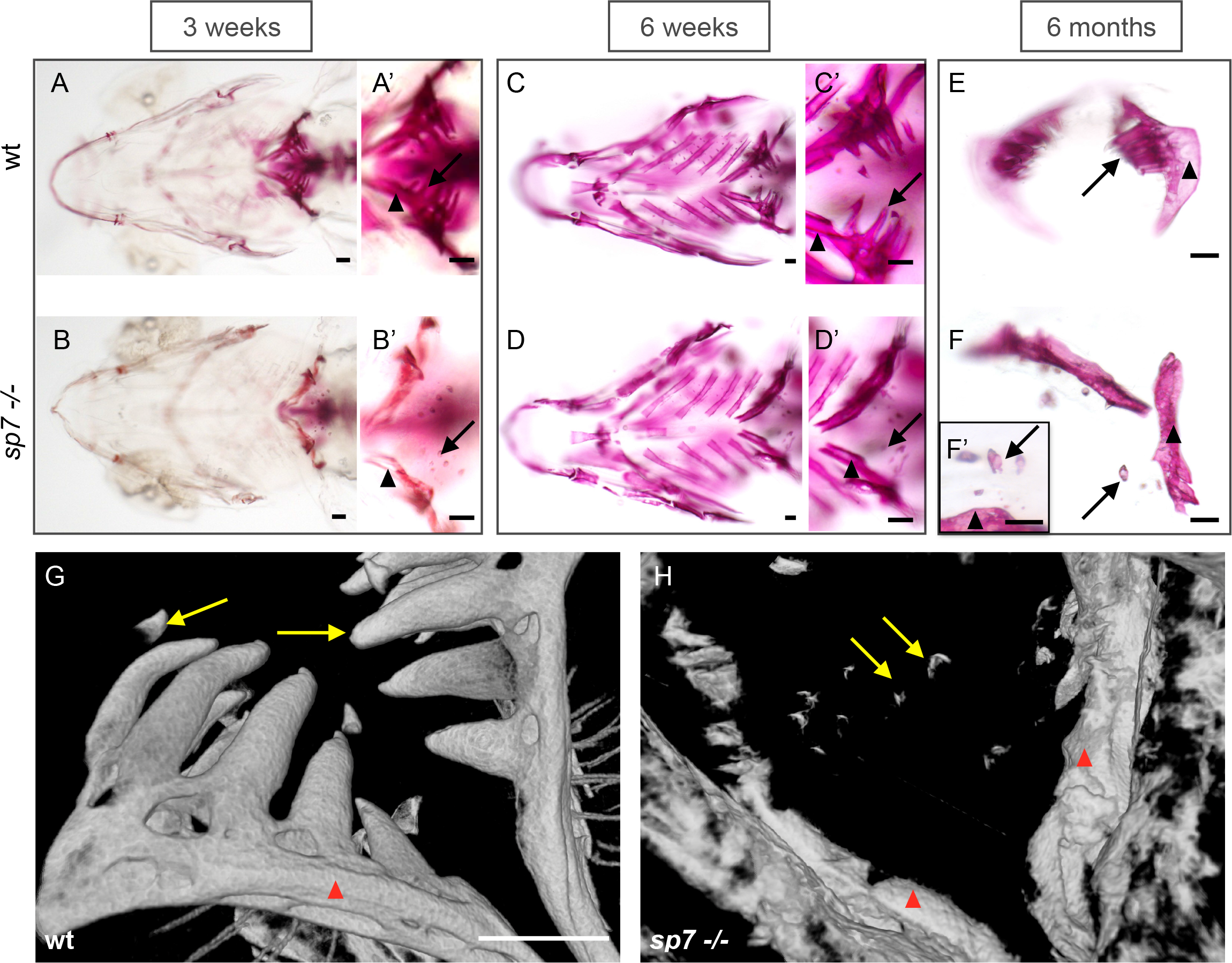
Defective tooth formation and lack of attachment in sp7 mutants observed throughout life. Alizarin Red S staining to show mineralization of the zebrafish pharyngeal teeth and fifth ceratobranchial (pharyngeal jaw) (A-F). A, A’, B, B’) In comparison to the wt, the fifth ceratobranchial (arrowheads) is poorly mineralized in mutants at 3 weeks post-fertilization, with only the extreme tip of the teeth stained (arrows). C, C’, D, D’) At 6 weeks, mineralization of the fifth ceratobranchial has increased in the mutants, but teeth remain as before. E, F, F’) *sp7* mutant pharyngeal bone displays a more narrow and thinner morphology, and teeth do not have a mineralized attachment to the bone. G, H) Volume rendering pictures of micro-computed tomography (µCT) of zebrafish (>1.5 year) to show pharyngeal bones (arrowheads) and teeth (arrows) in wt (G) and *sp7* mutant (H). Note the shape of the pharyngeal bones in the *sp7* mutant and the absence of any bone of attachment. Scale bars represent 100 µm.

### Differentiated odontoblasts express sp7 in wt zebrafish

To identify which cells express *sp7* during odontogenesis, we used adult zebrafish carrying the transgene *Tg(sp7:mcherry)*, evaluating expression by whole mount imaging (Fig. 2A-C), 3D renders of confocal z-stacks (Fig. 2D-E) and immunohistochemistry assays (Fig. 2F-L). *sp7* expression was detected in the pulp cavity of some teeth, but not in others, and detected in cells paving the inner lining of the attachment bone (Fig. 2A-C, D-E, F-J). Cross sections of 3D images show that within the pulp cavity, cells expressing *sp7* were localized in proximity to the dentin. Moreover, *sp7* expression levels correlate with the degree of tooth maturity (Fig. 2D’, E’, F, G, I, J). Single osteoblasts expressing *sp7* were clearly observed lining the bone of the pharyngeal jaws (Fig. 2D, E). For cellular resolution, we performed immunohistochemistry on sections followed by a detailed comparison with sectioned wt material stained with toluidine blue. The data revealed that expression of *sp7* within the pulp is clearly confined to odontoblasts lining the dentin (Fig.1. F, G). Young functional teeth still containing polarized odontoblasts in the tooth tip (Borday-Birraux et al., 2006) displayed *sp7* immunoreactivity (Fig. 2F-H), while expression was completely lost in firmly attached functional teeth with flattened odontoblasts (Fig. 2I-L). *sp7* expression was not observed in the ameloblasts in any of the developmental stages of the tooth. Together, these data indicate that *sp7* is expressed in actively secreting, differentiated odontoblasts but is downregulated with further maturation of the odontoblasts.

**Fig 2.**
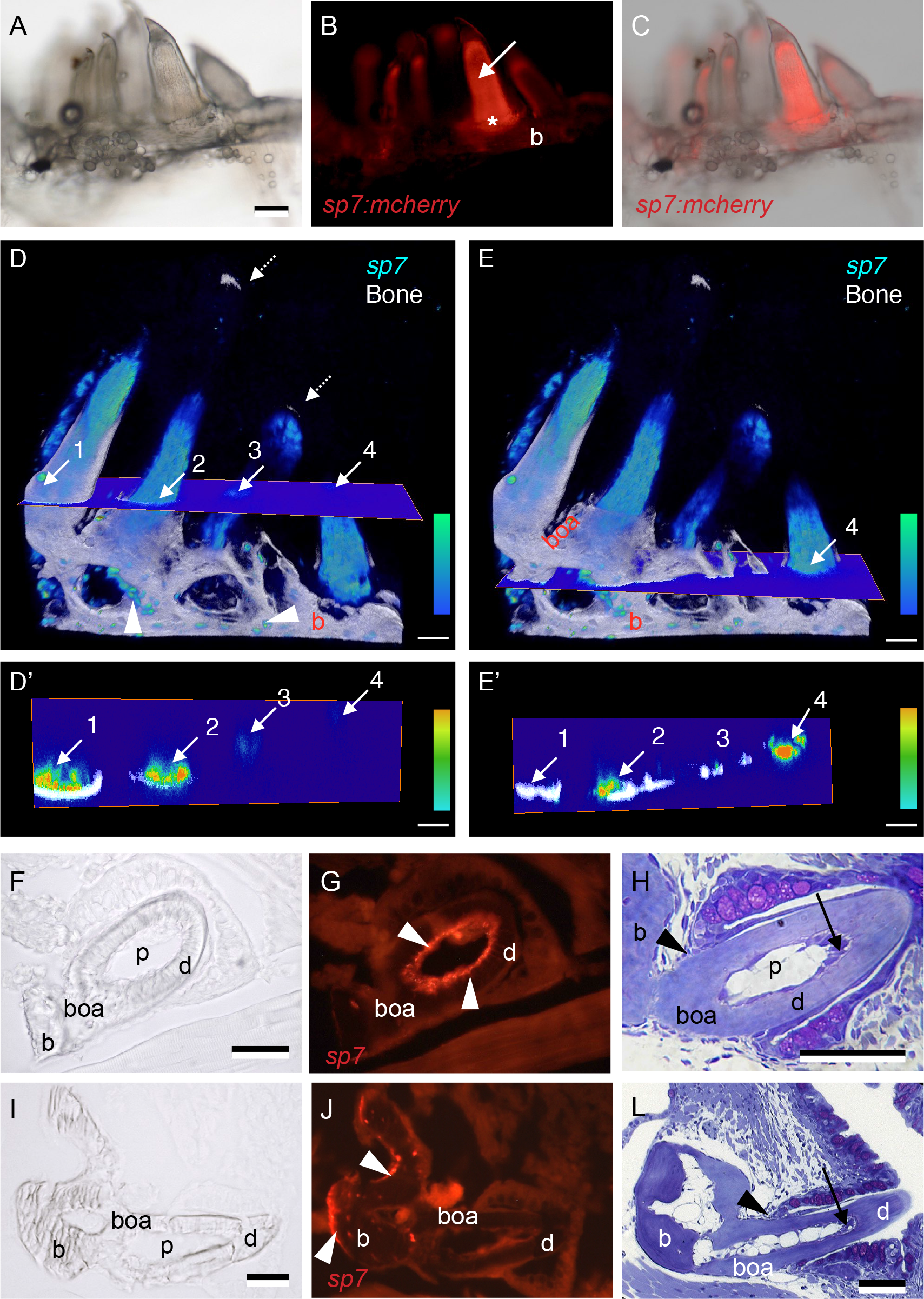
sp7 expression in the pharyngeal bones and teeth of zebrafish. Whole mount (A-C), 3D renders from confocal images (D-E’) and immunostained frozen sections (F-G, I-J) of teeth in juvenile *Tg(sp7:mcherry)* zebrafish (14 mm SL), compared with toluidine blue stained sections of similar-sized wt zebrafish teeth (H, L). A) Juvenile zebrafish pharyngeal bone seen under transmitted light, with the full complement of teeth present. B) Same sample shown under fluorescent light revealing *sp7* expression in the pulp (arrow), bone of attachment (asterisk) and underlying ceratobranchial bone (b). Note that other teeth show partial to no expression. C) Overlay between A and B; note that *sp7* expression varies according to the maturation of each tooth. D-E) 3D volume renders of z-stack from confocal microscopy images of *Tg(sp7:mcherry)*, level of *sp7* expression is color-coded as indicated by a color bar (bottom right), bone is coloured in white. Note that expression is stronger in some teeth but partial in others. Four teeth are indicated with numbers; arrows indicate where a cross section hits each tooth in two different planes (D and E). Isolated osteoblasts expressing *sp7* (arrowheads) are observed lining the pharyngeal bone (b), bone of attachment is clearly observed (boa). D’-E’) Corresponding cross section planes from D-E, one plane in a higher position (D’) than the other (E’). The same four teeth are indicated with arrows and numbers. Note that expression of *sp7* is stronger in proximity of the mineralized dentin (tooth 1 and 2). F-G) Cross section of a young functional tooth, immunostained for *sp7*, seen in transmitted (F) and fluorescent (G) light. Note odontoblasts (white arrowheads) expressing *sp7* in a linear arrangement juxtaposed to the dentin (d), continuing along the bone of attachment (boa). H) Toluidine blue stained plastic section of young functional tooth in a wt zebrafish. Note that odontoblasts at the tooth tip are still tall and polarized (black arrow). Black arrowhead indicates the level of the cervical loop and thus the limit between dentin (d) and bone of attachment (boa). I-J) Cross section of a mature functional tooth, immunostained for *sp7*, seen in transmitted (I) and fluorescent (J) light. Note lack of *sp7* expression in the pulp (p), while osteoblasts (white arrowheads) express *sp7* along the pharyngeal bone (b). L) Toluidine blue stained plastic section of a mature tooth in a wt zebrafish. Note that all odontoblasts have taken on a flattened shape, even in the tooth tip (black arrow). Black arrowhead indicates the level of the cervical loop and thus the limit between dentin (d) and bone of attachment (boa). Abbreviations: b, pharyngeal bone (fifth ceratobranchial); boa, bone of attachment; d, dentin; p, pulp. Scale bars represent 100 µm in (A-E’) and 50 µm in (F-L).

### Patterning and replacement of teeth is unaffected in sp7 mutants

To study the patterning of teeth in *sp7* mutants, crosses with *Tg(RUNX2:egfp)* were carried out to obtain mutants carrying the transgene labelling early odondoblasts, which permits tooth identification even without mineralization. A wild type pattern of teeth was observed in about 75% of mutant fish analysed (15 out of a total of 20 fish) with teeth arranged in rows running antero-posteriorly from medial to lateral (Fig. 3A-A’’ and B-B’’). The result was clearly confirmed with serial semi-thin sections of 6 months old mutants, showing eleven positions present on either body side. Thus, lack of *sp7*, and of the attachment structures that require *sp7*, does not appear to significantly affect the spatiotemporal patterning of the teeth. Interestingly, we found some teeth pointing towards random positions and away from the pharyngeal cavity (Fig. 3B’ and B’’) and some tooth pulps with irregular shape.

**Fig 3.**
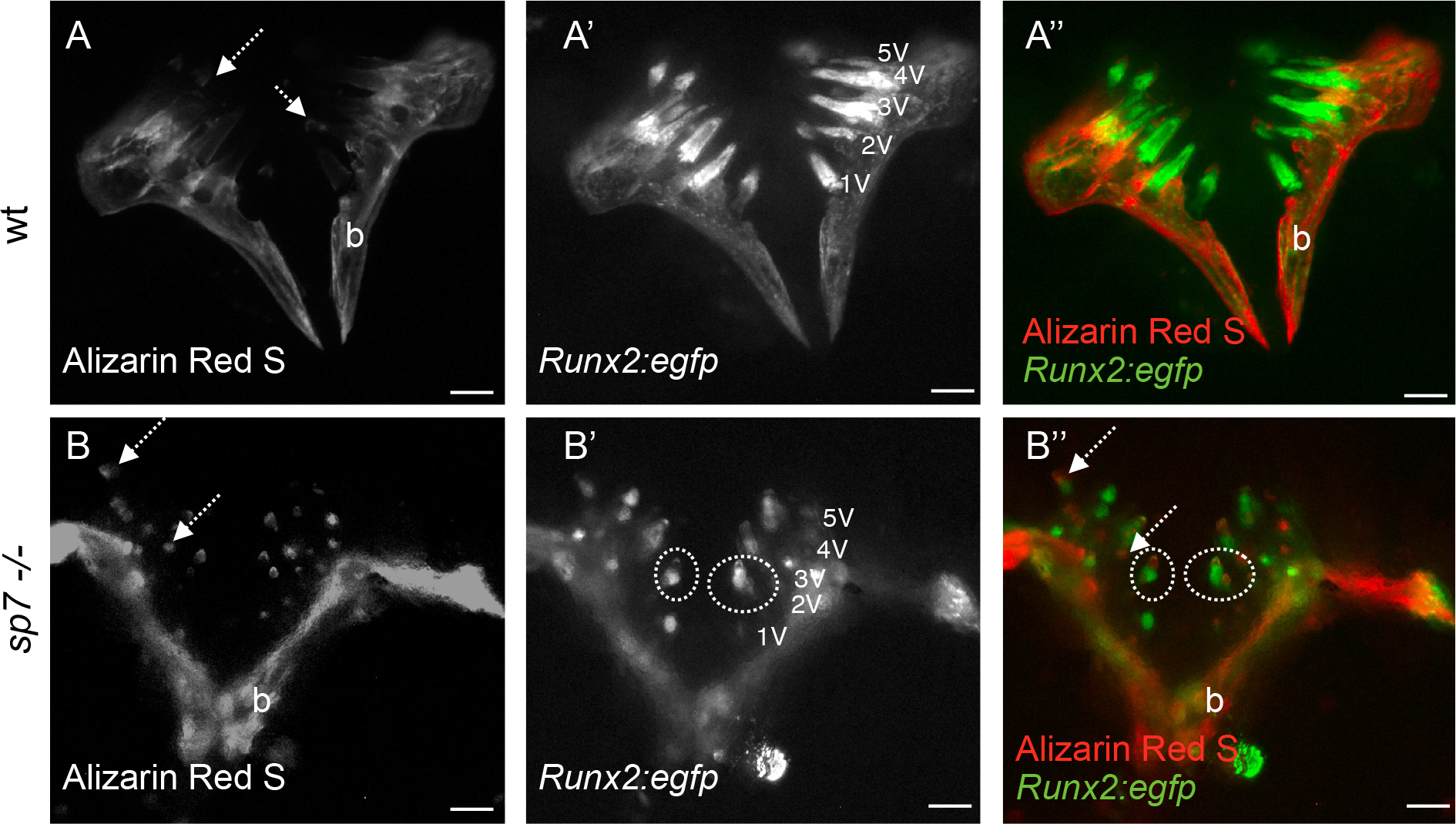
Tooth pattern in sp7 mutant is unaltered. A-B) wt and *sp7* mutant carrying *Tg(RUNX2:egfp)* and live stained for Alizarin Red S followed by dissection of pharyngeal bone and observed under a stereomicroscope and fluorescent light. Alizarin Red S stains calcium deposition in mineralized tissues. Note tooth tips strongly stained in mutants (dashed arrows) (A,B). Despite the number of teeth in *sp7* mutants being seemingly higher, it is still within the range that can be expected: each pharyngeal jaw can show as many as (maximum) 22 teeth (11 functional teeth, 11 replacement teeth). A’, B’) *Tg(RUNX2:egfp)* showing early odontoblasts and osteoblasts in pulp and bone respectively. Ventral tooth positions are indicated. Some tooth pulps display different shapes (dashed circles). Merged pictures with Alizarin Red S and *RUNX2:egfp* are shown (A’’, B’’). Scale bars represent 250 µm.

### Tooth development is severely disturbed in sp7 mutants

Although the number and position of teeth appeared to be normal, teeth displayed anomalous dentinogenesis (Fig. 4). An enamel organ was present as differentiation occurred into an inner and an outer dental epithelium (Fig. 4B, C compare with wt, A). The inner dental epithelium differentiated into polarised ameloblasts, and even ruffled-bordered ameloblasts. A cap of enameloid was laid down. Subsequently, a small amount of dentin was deposited. Yet, the odontoblasts never took on a cylindrical or polarised shape, and in none of the teeth could a layer of dentin be observed thicker than 6-8 µm, compared to 15-20 µm of thickness in wt teeth (Fig. 4B-C, F). Different from the wt, traces of organic material persisted in the mutant enameloid (Fig. 4C). Unlike in wt teeth, in which the pulp becomes extremely loosely organized with few cellular elements (Fig. 4D and H), the pulp cavity in the mutants remained highly cellular, filled with densely packed mesenchymal cells (Fig. 4B-C, F). We also observed tooth germs displaying an irregular orientation, with the tip pointing laterally rather than medially, or even turned away from the epithelial surface (Fig. 4E, compare to D). Thus, in the absence of *sp7*, development is arrested in early dentinogenesis.

**Fig 4.**
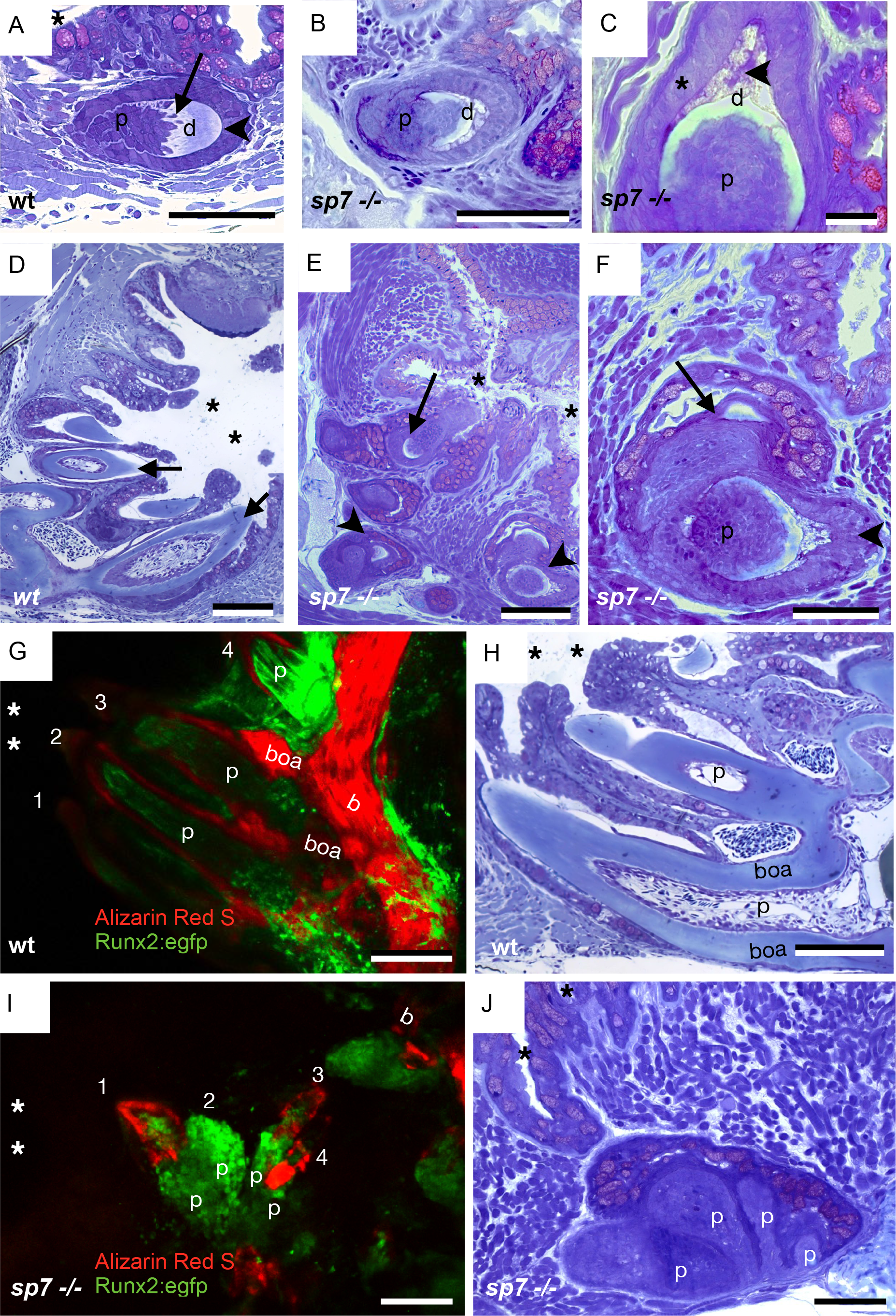
Tooth abnormalities in the sp7 mutant. Toluidine blue stained 2 µm sections of teeth in 6 months old wt (A, D and H) and *sp7* mutant (B-C, E-F and J) zebrafish. Confocal imaging of teeth in 5 months wt and *sp7* -/- zebrafish (G and I, respectively). A) Replacement tooth of a wt, with enameloid (arrowhead) and dentin (d) containing tubules (arrow), with predecessor tooth (asterisk) partly visible, localized in proximity of the pharyngeal epithelium. B) Overview of a defective tooth in the *sp7* mutant with a thin layer of dentin (d) and highly cellular pulp (p). C) Magnification of a *sp7* mutant tooth. Note small amount of dentin (d) and enameloid with traces of organic matter (arrowhead) but well organized enamel organ with polarized ameloblasts (asterisk). Note densely cellular pulp (p). D-E) Lower magnification picture to show the pharyngeal cavity (asterisks) and tooth arrangement in wt (D) and *sp7* mutant (E). In wt, teeth are oriented with tooth tips (arrows) pointing to the pharyngeal cavity (asterisks), while in the *sp7* mutant some teeth have an abnormal orientation with their tip (arrow) turned away from the pharyngeal cavity. Pairs of teeth can also be observed (arrowheads). F) A pair of tooth germs (predecessor, arrow and successor, arrowhead). These germs have an orientation relative to each other much as in the wt (compare with A). Note that the densely cellular pulps of both germs are interconnected without sharp boundary. G-J) Confocal imaging (G, I) and histological sections (H, J) of similar teeth to show pulps in wt (G, H) and *sp7* mutant (I, J). G, I). wt (G) and *sp7* mutant (I) carrying (Tg(*RUNX2*:egfp) and stained for Alizarin Red S, imaged using confocal microscopy followed by 3D projection. The pharyngeal cavity region (asterisks), pharyngeal bone (b) and bone of attachment (boa, in wt only) are labelled. Numbers label the different teeth. Note that the bone of attachment delimits each pulp (p) in the wt (G) while in *sp7* mutants (I) the lack of bones of attachment leads to abnormally connected pulps (p). H, J) Toluidine blue stained 2 µm cross section of pulps (p) in wt (H) and multi-pulps (p) connected in the *sp7* mutant (J). The pharyngeal cavity region is indicated by asterisks and the bone of attachment (boa) is labelled in the wt. Scale bars A, B, F and J = 50 µm; C = 20 µm; D, E, G, H and I= 100 µm.

### Tooth replacement continues despite the lack of tooth attachment

Serial cross sections of sp7 mutant fish revealed that teeth usually occurred in pairs (Fig. 4E, F). One, larger, tooth germ was often observed to have its tip protruding through the epithelial lining of the pharynx (hence erupted). The other tooth germ of a pair was seen to be located ventral to the larger germ, attached to the base of the pharyngeal crypt, corresponding to the position of a replacement (= successor) tooth in wt. The simultaneous presence of both teeth in *sp7* mutant fish shows that tooth replacement is ongoing. However, unlike in the wt, the pulp cavity of both teeth was observed to be disorganised and often abnormally connected (Fig. 4F). For the purpose of 3D visualization of pulp connectivity between multiple teeth, *sp7* mutants carrying Tg(RUNX2:egfp) were stained with Alizarin Red S and imaged using confocal microscopy followed by 3D projections. In wt, the tooth pulp is delimited by the bone of attachment (Fig. 4G, H). In contrast, in *sp7* mutants a multi-pulp structure, in which the pulp of predecessor and successor teeth are connected through *Runx2* positive cells, could be visualized with 3D projections (Fig. 4I) and in a single plane (Fig. 4J).

### sp7 mutants display normal osteoclast activity

In wt tooth cycling, resorption by osteoclasts, respectively odontoclasts, has two main functions: the resorption of worn functional teeth, and the preparation of the bone surface for the attachment of replacement teeth (Witten and Huysseune, 2009). Given the importance of osteoclasts for tooth cycling it was investigated whether osteoclasts are involved in resorption of predecessor teeth. We observed that osteoclastic activity was involved in resorption of the predecessor tooth (Fig. 5C), however this was less intense than during wt tooth replacement (compare with Fig. 5A). No increased resorption was observed that could explain the lack of tooth attachment. Remodelling of the ceratobranchial bone opposite growing teeth was observed in mutants at levels comparable to that seen in wt (compare Fig. 5B and D). Many osteoclasts appeared to be flat and mononucleated; typical for teleost osteoclasts (Witten and Huysseune, 2009).

**Fig 5.**
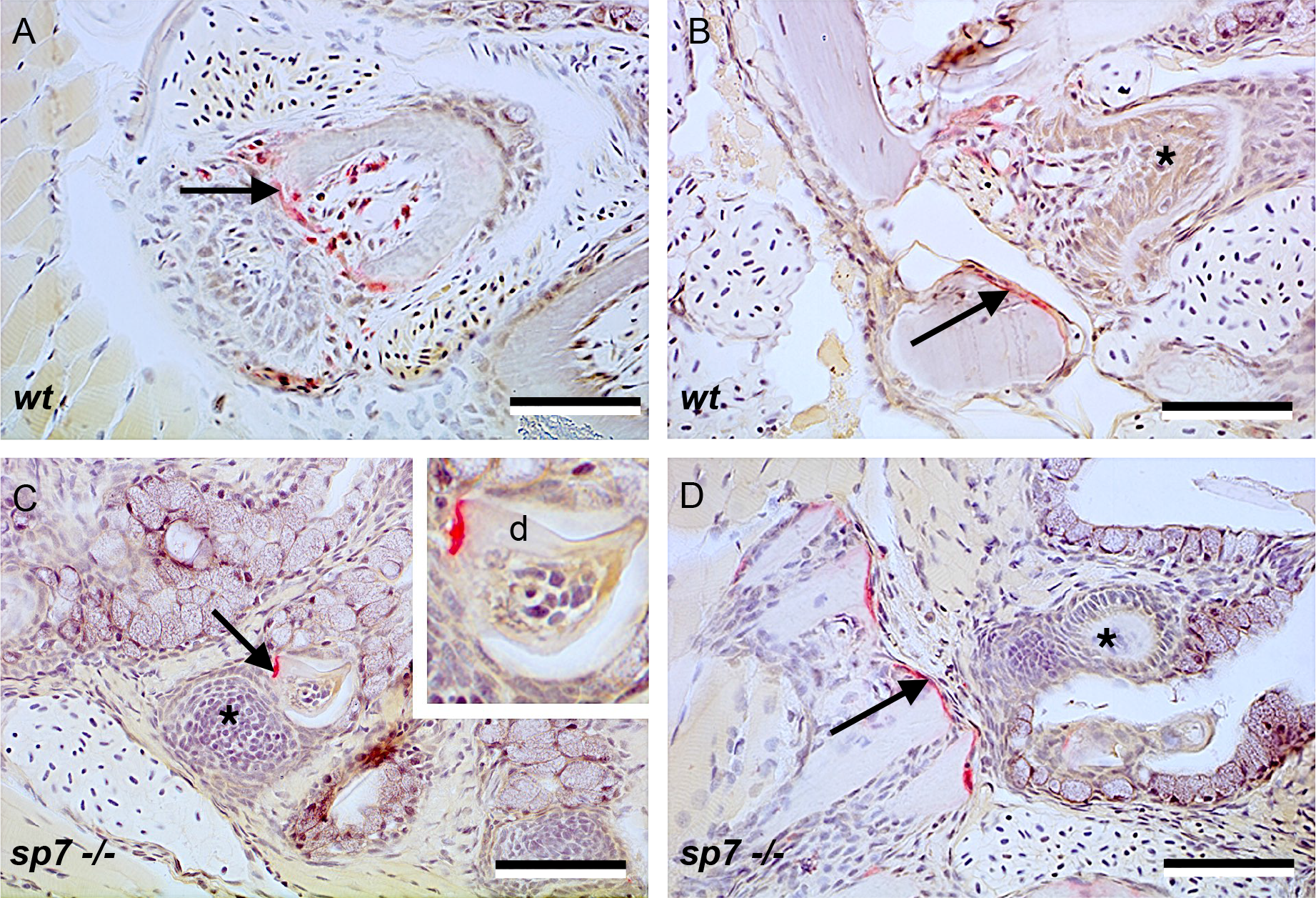
Osteoclastic activity revealed by TRAP staining. TRAP activity (red staining, arrows) in the predecessor tooth in wt (A) and *sp7* mutant (C) zebrafish, likely elicited by growth of a replacement tooth (asterisk in C). Inset in C shows magnification of TRAP activity involved in removal of the dentin (d). Note small size of the mutant teeth compared to the wt (A). TRAP activity (red staining, arrows) involved in remodelling of the ceratobranchial bone opposite a developing tooth (asterisk) is not different between wt (B) and mutant (D) fish. Scale bars = 50 µm.

## Discussion and Conclusions

Our data demonstrate that *sp7* is expressed in differentiated odontoblasts involved in dentin deposition, and in osteoblasts depositing the cylinder of attachment bone below the dentin base. The gene is downregulated in both cell types once the tooth has fully matured. This is once attachment bone has been fully laid down, dentinogenesis has been completed, and odontoblasts take on a flattened – interpreted as “resting” - phenotype. Such a downregulation is also seen in maturing odontoblasts of mice (Miyazaki et al., 2017). Overall, expression data are to a large extent consistent with data for mice (Chen et al., 2009) although the expression appears to be more mesenchyme-specific in zebrafish. Chen et al. observed *Sp7* expression in mice in mesenchymal condensates giving rise to bones and teeth at E12, and in mesenchymal cells in alveolar bone, dental papilla and follicle in cap stage teeth (E14) (Chen et al., 2009). At bell stage (E16) *Sp7* was expressed in odontoblasts and dental pulp cells (the signal persisting here until PN14) but also in ameloblasts. At no time did we observe expression in cells located in the zebrafish pulp cavity, other than odontoblasts, nor in ameloblasts.

Along with its expression in root odontoblasts of rats, *Sp7* is also expressed in cementoblasts, but not cementocytes (Hirata et al., 2009). Cementum is absent in teleosts; expression in zebrafish is nevertheless seen in osteoblasts lining the attachment bone, constituting the only structure securing attachment of the tooth to the underlying bone.

The defective dentinogenesis and the lack of attachment bone deposition observed in the zebrafish *sp7* mutants is consistent with the expression described above. The data also correspond to what has been recently reported for sp7-defective medaka *(Oryzias latipes)* (Yu et al., 2017). Medaka *sp7* mutants show a more severe bone phenotype than zebrafish *sp7* mutants. Medaka *sp7* mutants mineralize the vertebral body anlagen in the notochord sheath but fail to form the neural and haemal arches of the vertebral column. Moreover, medaka homozygous mutants do not survive to adulthood. As medaka has numerous oral and pharyngeal teeth and tooth patterning is different from zebrafish, direct comparisons are difficult. However, Yu et al. reported a smaller tooth size in medaka mutants due to a reduction in dentin deposition and the absence of attachment bone (Yu et al., 2017). Both in zebrafish and medaka, this phenotype is in line with a role for the gene in mammals for the formation of root, but not crown, dentin and of cellular cementum (Cao et al., 2012; Zhang et al., 2015). In many teleosts, the dentin base is ankylosed to the underlying bone via a short cylinder of attachment bone. *sp7* mutants are defective in formation of an attachment structure, suggesting that, despite fundamental differences between attachment bone and cementum, some commonality exists in both ways of ankylosis. The evolutionary history of attachment bone in actinopterygians remains unresolved, but its close developmental relationship to the tooth unit is evident (Sire and Huysseune, 2003). The simultaneous presence of transcripts in odontoblasts and in osteoblasts that line the attachment bone, and the defect both in dentin and attachment bone formation elicited by the absence of *sp7*, indeed suggests common regulatory mechanisms for both tissues. The lack of a normal dentin shaft and attachment bone can explain why the pulp of the functional tooth, and of its successor, remain connected. The apparently undermineralized enameloid seen in *sp7* mutants, along with traces of persisting organic matrix, can be ascribed to the defective function of the odontoblasts, given the involvement of these cells in the production of the enameloid (Sasagawa et al., 2009).The patterning of the teeth in zebrafish (three rows, with 5, 4 and 2 teeth respectively on each body side) is established with the first-generation teeth and is maintained into adulthood probably through a local control mechanism (Borday-Birraux et al., 2006). This pattern appears to be maintained at least in mildly affected mutants. Pulp connection between predecessor and replacement tooth was often observed. In such cases the replacement tooth is forced to form under a different orientation; such change can disturb the orientation of the following successional tooth and so on, overtime leading to a misplacement and poor orientation of teeth. The lack of attachment of the teeth to the underlying bone might play a role in the way how the pulps are connected. However, it does not prevent eruption of some of the teeth, as revealed by the fact that some tooth tips pierce through the epithelium. This is not surprising, given that eruption of zebrafish teeth has been observed to be triggered by epithelial remodelling and not by growth or movement of the tooth (Huysseune and Sire, 2004).

Given that *sp7* knockout mice die at birth (Nakashima et al., 2002) and wt mice have only one tooth generation, mice are not an appropriate model to study the role of the gene in tooth replacement. Contrary to earlier insights, the current data reveal that neither attachment, nor eruption, is required to stimulate the initiation of the successor tooth. It was formerly suggested that movement of the epithelial layers could displace the putative stem cell niche thus stimulating the formation of a new tooth bud in wt zebrafish (Huysseune and Thesleff, 2004). Here, we show that defective differentiation of the tooth does not prevent a successional tooth to be formed.

In conclusion, our data on *sp7* expression and function during zebrafish odontogenesis are in line with its role in root formation in mammals, in that dentinogenesis is started but not achieved, and attachment fails. Neither attachment nor eruption is required for initiation of the new, successional tooth. These data further emphasize the validity of zebrafish as a useful model to study tooth development and regeneration.

## Ethics

Experiments were approved by the local ethics committee and were granted a UK Home Office project licence.

## Author contributions

E Kague, A Huysseune contributed to conception, design of the experiments and drafting of the manuscript. PE Witten contributed to data collection, analysis, draft of manuscript, and revision of the manuscript. S Fisher, T Lubiana, C Lovaglio and M Soenens contributed to data acquisition. C Hammond, K Robson Brown (µCT data acquisition) and MR Passos-Bueno contributed to data acquisition and revision of manuscript.

## Acknowledgements/Funding

C Hammond and E Kague were funded by Arthritis Research UK (19476, 21211). We acknowledge financial support from CNPQ (National Council for Scientific and Technological Development, Brazil) (process 407034/2013-7) and FAPESP (São Paulo Research Foundation, Brazil)/CEPID (process 2013/08028-1). AH gratefully acknowledges a grant from the BOF-Ghent University, project number 01J05815. The authors declare no potential conflicts of interest with respect to the authorship and/or publication of this article.

